# Strigolactone signaling regulates corm development through SPL15-mediated hormonal crosstalk in banana

**DOI:** 10.64898/2026.03.12.711407

**Authors:** Fang Long, Ming Zhao, Peng Wu, Yu Zhou, Xiang Xiang, Tian-li Mo, Xiangyu Hu

## Abstract

Strigolactones (SLs) are an important class of plant hormones that play crucial roles in regulating plant branching, root architecture, and organ development. However, the regulatory mechanisms underlying the crosstalk between SLs and other plant hormones remain largely unclear, particularly regarding the key regulatory genes that integrate and coordinate multiple hormonal signaling pathways. In this study, secondary cup seedlings of the Pisang Awak banana cultivar ‘Yufen 6’ at the eight-leaf stage were used as experimental materials. The roots were treated with a nutrient solution containing 30 μmol/L exogenous SLs, while a nutrient solution supplemented with water served as the control. Tissues near the corm growth point were collected at 0, 15, 30, 60, 90, and 120 days after treatment to measure corm weight, height, and diameter, and transcriptome sequencing was performed using the collected tissues. Differentially expressed genes (DEGs) at different treatment stages were identified, followed by Gene Ontology (GO) annotation and Kyoto Encyclopedia of Genes and Genomes (KEGG) pathway enrichment analyses to systematically investigate the crosstalk between SLs and endogenous hormone metabolism and signaling during corm development in Pisang Awak banana.

The results showed that SL treatment significantly inhibited the weight, height, and diameter of the corm. The regulatory effect of SLs on Pisang Awak banana corm development exhibited a clear temporal dynamic pattern, representing a gradual accumulation process that ultimately triggers key developmental transitions. The highest number of DEGs was detected at 15 days after treatment, including 3943 upregulated genes and 3704 downregulated genes, indicating that this stage represents a critical phase for SL response initiation. GO enrichment analysis revealed that the DEGs were mainly involved in metabolic processes, biological regulation, response to stimulus, and regulation of biological processes. KEGG pathway analysis indicated that these DEGs were significantly enriched in pathways related to plant hormone signal transduction, starch and sucrose metabolism, and secondary metabolite biosynthesis. Further analysis revealed that the crosstalk between SLs and multiple hormone metabolic and signaling pathways is mediated by the SPL15 gene, involving auxin (IAA), cytokinin (CTK), abscisic acid (ABA), brassinosteroids (BRs), gibberellins (GA), and jasmonic acid (JA) pathways. This study reveals the molecular mechanism by which SLs regulate Pisang Awak banana corm development through SPL15-mediated integration of multiple hormonal signals, providing new insights into the role of SLs in regulating the development of underground organs in banana.

---

The corm of banana (*Musa* spp.) is a modified stem characterized by shortened internodes and a swollen spherical structure, commonly referred to as the “banana corm” ^1, 2^. It serves as an important organ for nutrient storage and sucker formation. Banana suckers originate from axillary buds located at the nodes near the base of the corm ^3^. The development of suckers essentially represents the formation of lateral branches. During growth, suckers compete with the mother plant for nutrients, which can significantly influence the production cycle and yield of banana ^4^. In agricultural practice, farmers often need to invest considerable labor and resources to remove excess suckers to maintain optimal plant growth and productivity. Therefore, understanding the regulatory mechanisms underlying sucker formation and the development of the banana corm is of great significance for improving cultivation management and enhancing production efficiency.

Strigolactones (SLs) are a class of plant hormones discovered in recent years. They are mainly synthesized in roots from carotenoid-derived metabolites and subsequently transported to the aerial parts of plants ^5^. Previous studies have demonstrated that SLs play crucial roles in regulating plant branching, lateral bud growth, stem elongation, the formation of adventitious and lateral roots, as well as root hair development, and they are also involved in plant responses to various environmental stresses ^6^. Increasing evidence indicates that SLs do not function independently; instead, they interact with multiple plant hormones to form a complex multilayered regulatory network that coordinates plant growth and development across different organs and developmental stages.

Previous studies have shown that SLs regulate lateral bud development by modulating the metabolism and signaling pathways of other plant hormones. For instance, in rice, SLs suppress branching by inhibiting the expression of the cytokinin (CTK) biosynthetic gene IPT while promoting the expression of cytokinin oxidase/dehydrogenase genes (CKX), thereby reducing CTK levels in lateral buds ^7^. In Arabidopsis, the SL signaling pathway activates the transcription factor BRC1 (BRANCHED1), which subsequently induces the expression of HB40 and further upregulates the abscisic acid (ABA) biosynthesis gene NCED3, leading to increased ABA accumulation in lateral buds and inhibition of bud outgrowth ^8^. In addition, SLs can repress the expression of auxin (IAA) biosynthesis genes such as YUCCA, thereby reducing IAA levels and polar auxin transport in lateral buds and ultimately inhibiting bud elongation ^9^. These studies suggest that SLs regulate plant branching through interactions with multiple hormonal pathways.

However, although previous studies have reported that SLs can regulate key genes in other hormone pathways—such as the CKX genes in the CTK metabolic pathway or genes involved in auxin biosynthesis—the molecular mechanisms by which SL signaling interacts with these hormonal pathways remain largely unclear. In particular, how SL signaling integrates and coordinates multiple hormone pathways through specific transcriptional regulators to achieve hormonal crosstalk during plant development has rarely been elucidated.

To date, studies on SL-mediated regulation of plant growth and development have mainly focused on model plants and major crops such as Arabidopsis ^10^, pea ^11^ and rice ^7^, primarily investigating lateral bud and root development. In contrast, the interaction mechanisms between SLs and other plant hormones in banana, an important tropical fruit crop, remain largely unknown. The banana corm, which serves as the central organ for sucker formation and nutrient storage, undergoes development regulated by multiple hormone metabolic and signaling pathways. However, the molecular mechanisms underlying SL-mediated regulation of banana corm development are still poorly understood.

In this study, the Pisang Awak banana cultivar ‘Yufen 6’ was used as the experimental material. Plants were treated with the synthetic SL analog rac-GR24, and corm tissues at different treatment stages were collected for transcriptome sequencing. By comparing the transcriptomic profiles between treated and control samples at multiple time points, we systematically analyzed the expression changes of genes involved in the metabolism and signaling pathways of several plant hormones, including cytokinin (CTK), auxin (IAA), abscisic acid (ABA), gibberellins (GAs), brassinosteroids (BRs), and jasmonic acid (JAs). Differentially expressed genes (DEGs) associated with SL-mediated regulation of banana corm development were identified, aiming to elucidate the crosstalk mechanisms between SLs and other plant hormones. This study provides new insights into the molecular mechanisms by which SLs regulate banana corm development and contributes to a better understanding of hormone interaction networks in banana.

## Methods

### Plant material

The experimental material used in this study was the Pisang Awak banana cultivar ‘Yufen 6’, developed by the authors’ team. Single suckers were collected from the mother plants in the Yufen 6 germplasm orchard at the Guangxi Academy of Agricultural Sciences after three consecutive sunny days and used for rapid propagation via tissue culture. After ten generations of in vitro propagation, the plantlets were grown into secondary cup seedlings. Healthy and uniform eight-leaf-stage secondary seedlings were selected for the experiment.

### Experimental design

Preliminary experiments indicated that treatment with 30 μmol/L rac-GR24 for 5 days had a notable effect on corm development. For this study, the roots of eight-leaf-stage plantlets were carefully washed and pre-cultured in Hoagland’s solution for 7 days. Rac-GR24 (Beijing Cool-Lab Technology Co., Ltd.) was dissolved in a small amount of acetone and kept in reserve. After pre-culture, uniform and healthy seedlings were retained, and the nutrient solution was replaced with Hoagland’s solution containing 30 μmol/L rac-GR24 for root shading hydroponic treatment. The control group received the same volume of Hoagland’s solution supplemented with water. The culture solution was renewed daily, and the treatment was applied for 5 consecutive days.

Following treatment (August 1, 2024), both treated and control plantlets were transplanted into field plots at the Yufen 6 base of Guangxi Academy of Agricultural Sciences under uniform soil conditions. The soil surface was covered with dry weeds, and a shade net with 60% light transmittance was installed at 1.0 m above the ground during daytime. Water and fertilizer management were standardized across all plots.

Corm tissues (including the growth point and with emerging suckers above the corm surface removed) were sampled at 0, 15, 30, 60, 90, and 120 days after the 5-day treatment. The samples were designated as SL0, SL15, SL30, SL60, SL90, and SL120 for the rac-GR24-treated group, and CK0, CK15, CK30, CK60, CK90, and CK120 for the control group. Corm height and diameter were measured, and tissues were cleaned, ground, and homogenized. Each treatment consisted of three biological replicates, with nine plants per replicate.

### RNA extraction, sequencing, and transcriptome analysis

Total RNA was extracted using the Trizol reagent kit (Invitrogen, Carlsbad, CA, USA), and RNA integrity was assessed with the Agilent 2100 Bioanalyzer (Agilent Technologies, Palo Alto, CA, USA). Samples with high-quality RNA were used for transcriptome sequencing. Non-strand-specific cDNA libraries were prepared using the NEBNext® Ultra™ II RNA Library Prep Kit (New England Biolabs, Ipswich, MA, USA) according to the manufacturer’s instructions. Sequencing was performed by Guangzhou Gidio Biotechnology Co., Ltd. on the Illumina HiSeq™ platform.

Raw sequencing reads were filtered to remove low-quality sequences and potential contaminants using fastp, generating clean reads for downstream analysis. Reads containing adapters, reads with more than 10% ambiguous bases (N), reads composed entirely of adenine (poly-A), and reads in which more than 50% of bases had a quality score ≤20 were removed. To eliminate ribosomal RNA sequences, clean reads were aligned to the species-specific rRNA database using Bowtie2, and reads mapped to rRNA were discarded. The remaining unmapped reads were aligned to the reference genome (GCA_004837865.1; https://doi.org/10.6084/m9.figshare.22716271.v9) using HISAT2. Transcripts were reconstructed using StringTie, and gene expression levels were quantified with RSEM. Differentially expressed genes (DEGs) were identified using DESeq2, with read counts normalized, statistical significance calculated, and multiple testing correction applied to control the false discovery rate (FDR). Genes with FDR < 0.05 and |log2 Fold Change| > 1 were considered significantly differentially expressed.

Functional annotation of DEGs was performed by mapping them to Gene Ontology (GO) terms, and enrichment was assessed using hypergeometric tests to identify GO terms significantly overrepresented in DEGs compared with the background. KEGG pathway enrichment analysis was further conducted to identify signaling and metabolic pathways significantly enriched in DEGs, providing insights into the potential molecular mechanisms of strigolactone-mediated regulation of banana corm development.

### Real-time quantitative PCR verification

Total RNA was reverse-transcribed into first-strand cDNA using the MonScript™ RTIII cDNA Synthesis Kit (Monad, USA) following a two-step qPCR protocol. The reverse transcription program consisted of incubation on ice for 5 min, 37 °C for 10 min, 55 °C for 15 min for cDNA synthesis, and 85 °C for 5 min to terminate the reaction. The synthesized cDNA was used as a template for subsequent quantitative real-time PCR (qRT-PCR).

Seven genes related to SL regulation and hormone crosstalk, including *SPL15* (ID: C4D60_Mb08t24630), *ARF15* (ID: BXH2_00010873), *ADH2* (ID: BXH2_00004643), *PIL13* (ID: C4D60_Mb05t02240), *CKX11* (ID: BXH1_00004522), *XTH22* (ID: BXH3_00010891), and *PYL3* (ID: C4D60_Mb10t15440), were selected for qRT-PCR validation, with *UBQ2* serving as the internal reference. Primers were designed using primer blast (Table 1) and synthesized by Shanghai Shenggong Bioengineering Technology Service Co., Ltd.

**Table 1.**
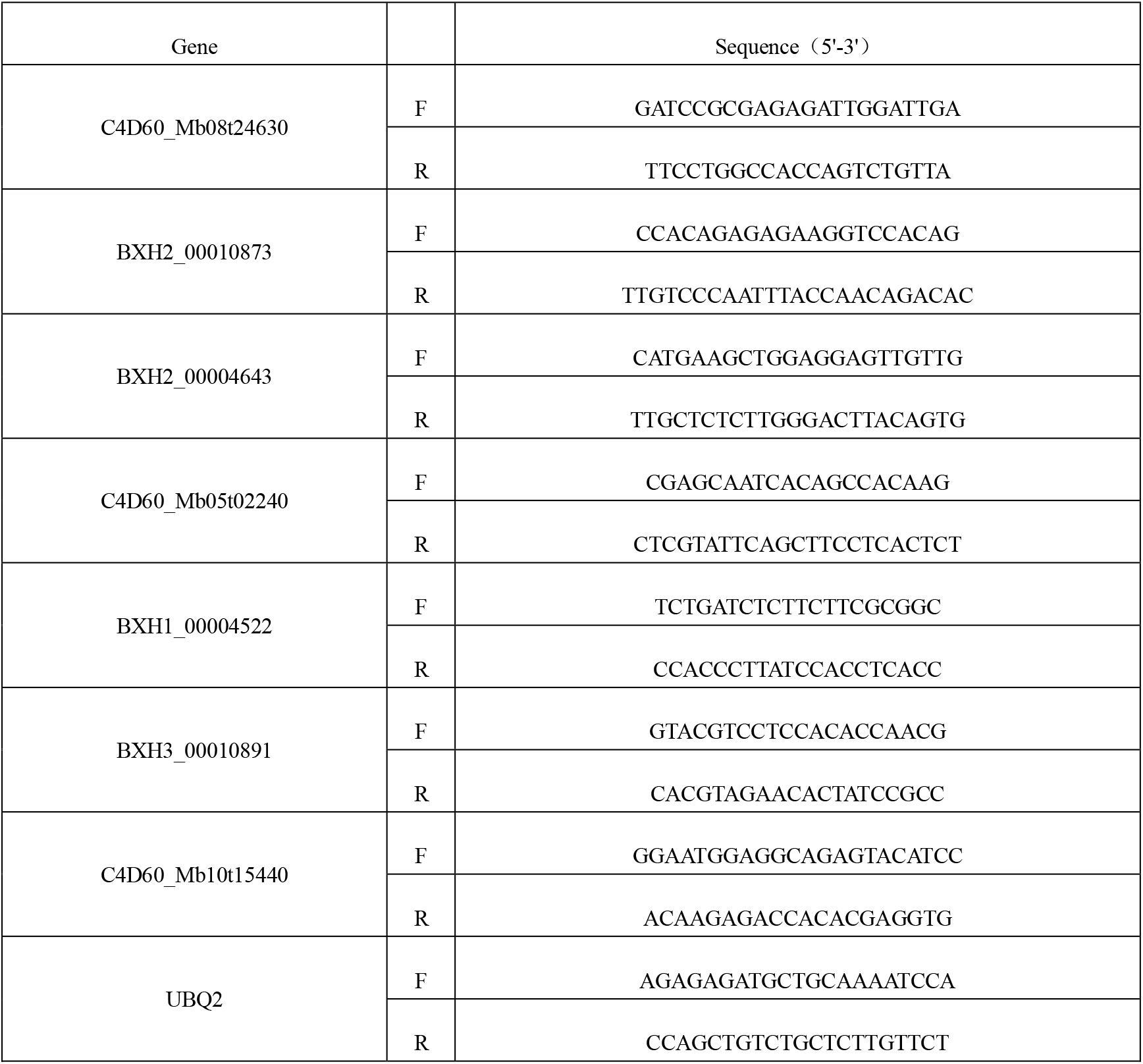
Real-time fluorescence quantitative PCR primer sequences.

The qPCR reaction mixture (10.0 μL) contained 5.0 μL of 2× SYBR Real-Time PCR Pre-mix, 0.05 μL of QN ROX Reference Dye, 0.7 μL each of forward and reverse primers, 1.0 μL of cDNA template, and RNase-free water to a final volume of 10.0 μL. The amplification program consisted of an initial denaturation at 95 °C for 5 min, followed by 40 cycles of 95 °C for 5 s and 60 °C for 30 s. Relative gene expression levels were calculated using the 2^^-ΔΔCt^ method ^12^.

### WGCNA analysis and cytoscape visualization

The gene expression matrix was first preprocessed in R. Genes were retained if they met the following criteria: at least half of the samples had TPM values greater than 1, and the median absolute deviation (MAD) exceeded the 50th percentile. Genes of interest, specifically those involved in hormone signaling pathways, were retained even if they did not meet these thresholds.

Weighted gene co-expression network analysis (WGCNA) was used to construct the co-expression network. The soft-thresholding power was determined based on the scale-free topology fit index to ensure the network approximated a scale-free topology, typically requiring R^2^ > 0.9; the selected power was 11. Pearson correlation coefficients were calculated between all gene pairs and transformed into an adjacency matrix using the formula adjacency = |cor|^power. The adjacency matrix was further converted into a topological overlap matrix (TOM) to reduce noise and improve network robustness. Hierarchical clustering based on TOM distance was performed, and modules were identified using the dynamic tree cut method with a minimum module size of 30 genes. Highly similar modules were merged according to the correlation of module eigengenes (MEs) with a merge cut height of 0.25. Module eigengenes were calculated and correlated with phenotypic data using Pearson correlation analysis to identify modules significantly associated with traits.

Candidate hub genes within each module were selected based on the criteria |module membership (MM)| > 0.8 and intramodular connectivity (K.in) ranking within the top 10% of all genes in the module. For network visualization, WGCNA was used to export node and edge files for Cytoscape, and edges with weights less than 0.1 were removed to enhance clarity.

## Results

### Effects of SLs treatment on corm phenotype

Strigolactone (SL) treatment significantly inhibited corm growth in Pisang Awak banana. From 15 days after treatment, corm weight, height, and diameter were all significantly lower than those of the control group (Figure 1).

**Fig. 1.**
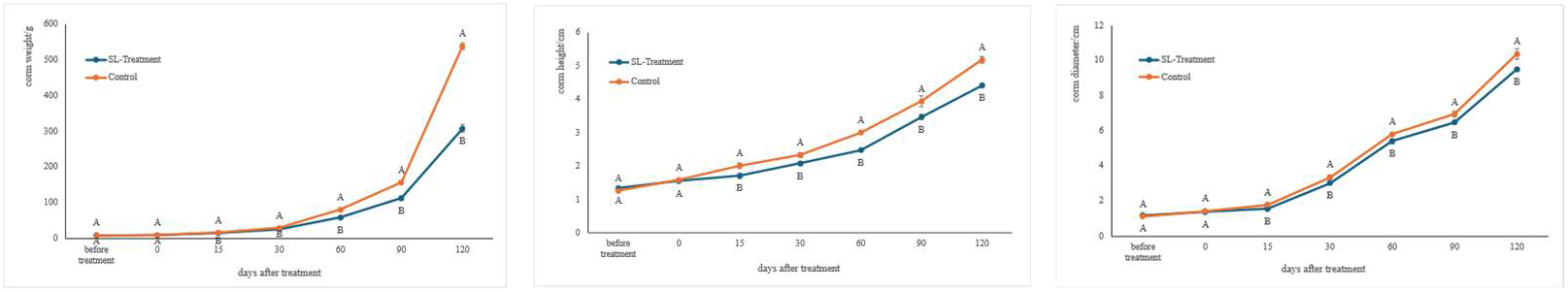
Changes of the growth and development of corm at different treatment time periods Different capital letters in the figure represent significant differences between the treatment and control in the same period (*P* < 0.05)

### Quality assessment of transcriptome sequencing data

Transcriptome sequencing generated 35,638,662 to 45,367,762 clean reads per library, with clean read proportions ranging from 99.54% to 99.80%, and mapping rates to the reference genome exceeding 94.96%, indicating high data quality and reliability (Figure 2).

**Fig. 2.**
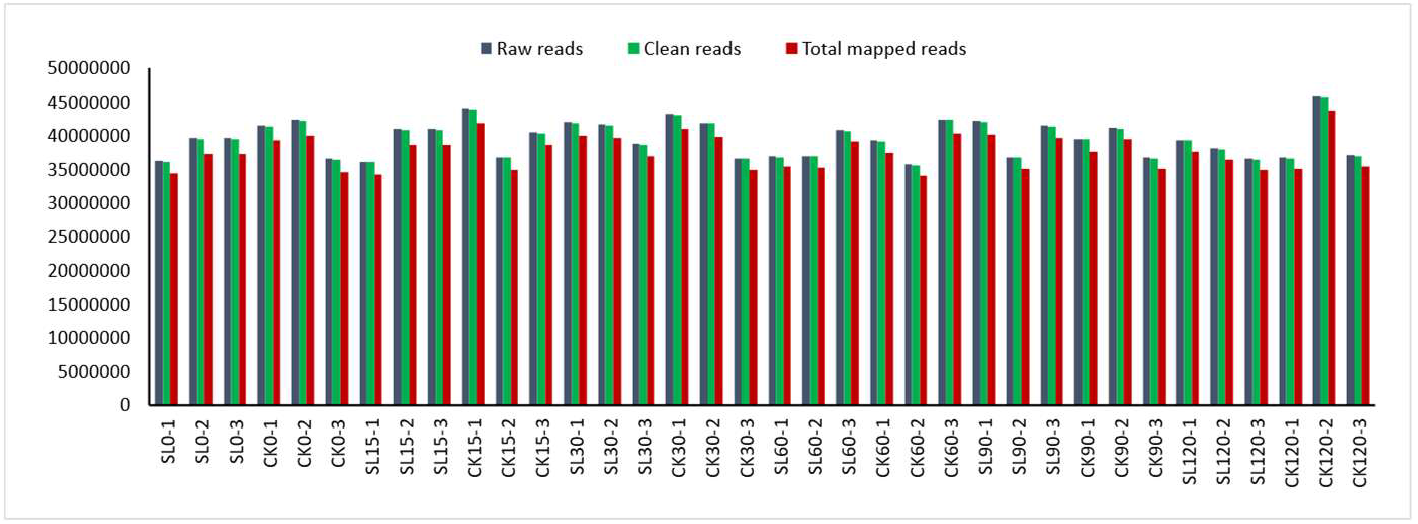
Overview of RNA seq data

### Dynamic changes of DEGs

Differential expression analysis revealed that the number of differentially expressed genes (DEGs) across developmental stages first increased and then decreased. Among the six sampling points, the second stage (15 days after treatment) exhibited the largest number of DEGs, with 3,943 upregulated and 3,704 downregulated genes, suggesting that this period represents a highly active phase of gene expression regulation in the corm. As development progressed, the number of DEGs gradually declined, reaching the lowest level at the sixth stage (120 days after treatment), with only 166 upregulated and 274 downregulated genes (Figure 3). These results indicate that SLs exert their regulatory effects primarily during the first 15 days after treatment, with their influence gradually diminishing thereafter.

**Fig. 3.**
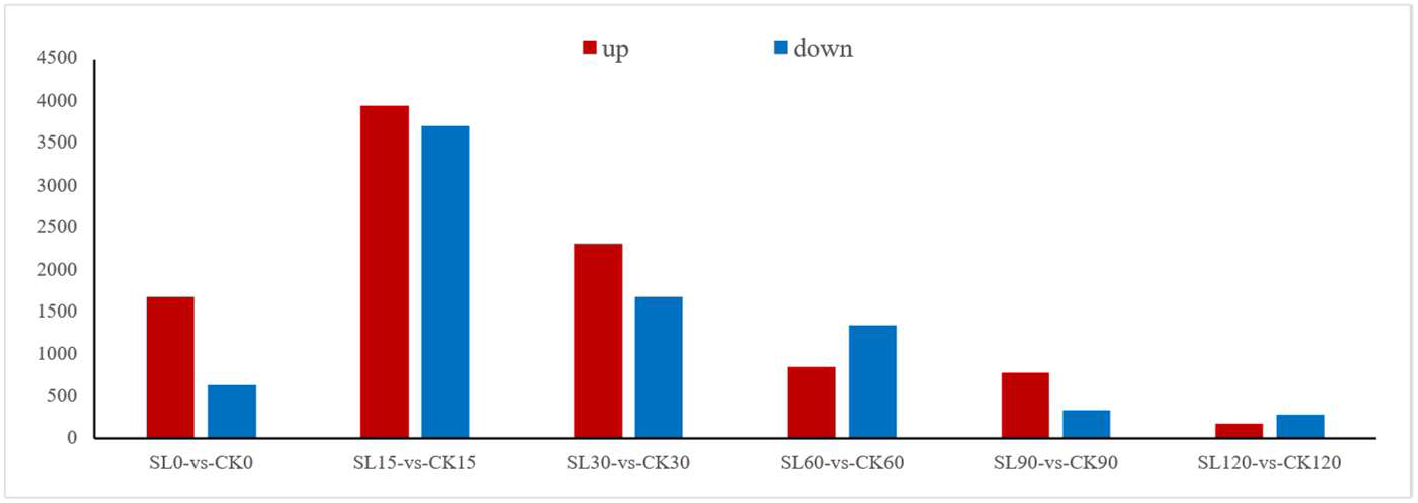
Number of DEGs between the treatment and control groups at different time

### GO and KEGG enrichment analysis of DEGs

GO enrichment analysis (Figure 4) revealed significant differences between SL0 and CK0 across multiple biological processes and molecular functions. Within biological processes, cellular process, metabolic process, biological regulation, response to stimulus, and hormone-related regulation were the most significantly enriched, indicating that SL treatment substantially affects cellular metabolism and hormone responses. Enrichment in signaling-related processes further suggested that SL-mediated regulation is closely associated with hormone signal transduction. For cellular components, genes were significantly enriched in cellular anatomical entities, membranes, and protein-containing complexes, implying potential involvement of structural proteins in regulation. Regarding molecular functions, catalytic activity and binding were most enriched, indicating that many DEGs encode enzymes, receptors, or transcription factors. Transcription regulator activity was notably upregulated, highlighting the importance of transcriptional control. Changes in antioxidant activity and transporter activity may relate to homeostasis maintenance and hormone transport. Although enrichment patterns were generally consistent across sampling time points, the number of DEGs peaked at 15 days and gradually declined thereafter.

**Fig. 4.**
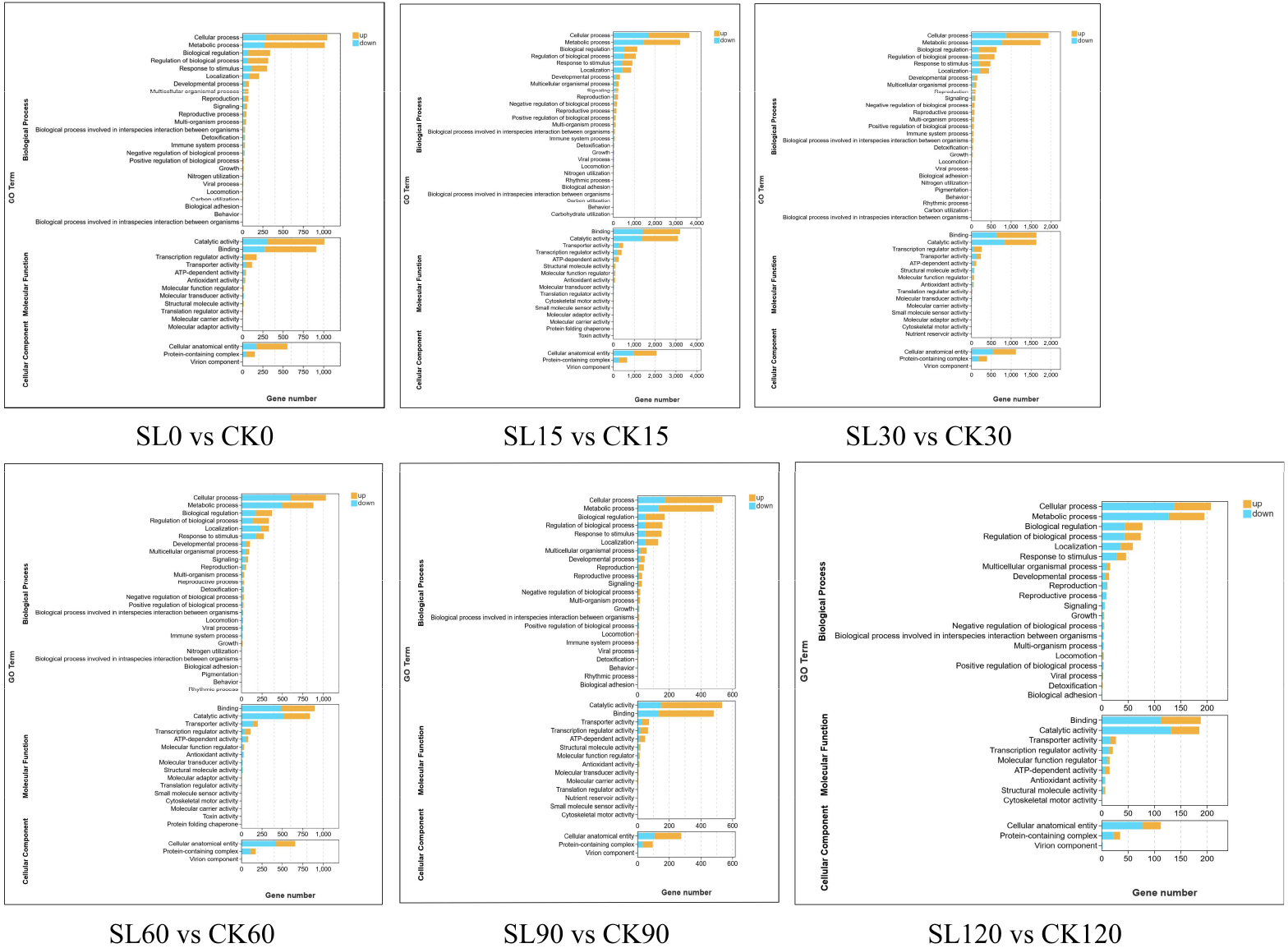
GO enrichment analysis of differentially expressed genes in different periods

KEGG pathway enrichment analysis (Figure 5) showed that at the early stage (0 days), DEGs were significantly enriched in plant hormone signal transduction, starch and sucrose metabolism, and secondary metabolite biosynthesis pathways, suggesting that hormone signaling and carbon metabolism respond rapidly to SL treatment. At 15 days, DEGs were significantly enriched in plant hormone signaling, nucleotide excision repair, and DNA replication pathways, indicating active cell division during this period, which may correspond to the peak phase of SL-mediated sucker initiation. In later stages, enriched pathways gradually included lipid metabolism (alpha-linolenic acid metabolism), amino acid metabolism (tryptophan metabolism), and secondary metabolite biosynthesis (phenylpropanoid biosynthesis), reflecting enhanced structural biosynthesis activity during the later developmental phases.

**Fig. 5.**
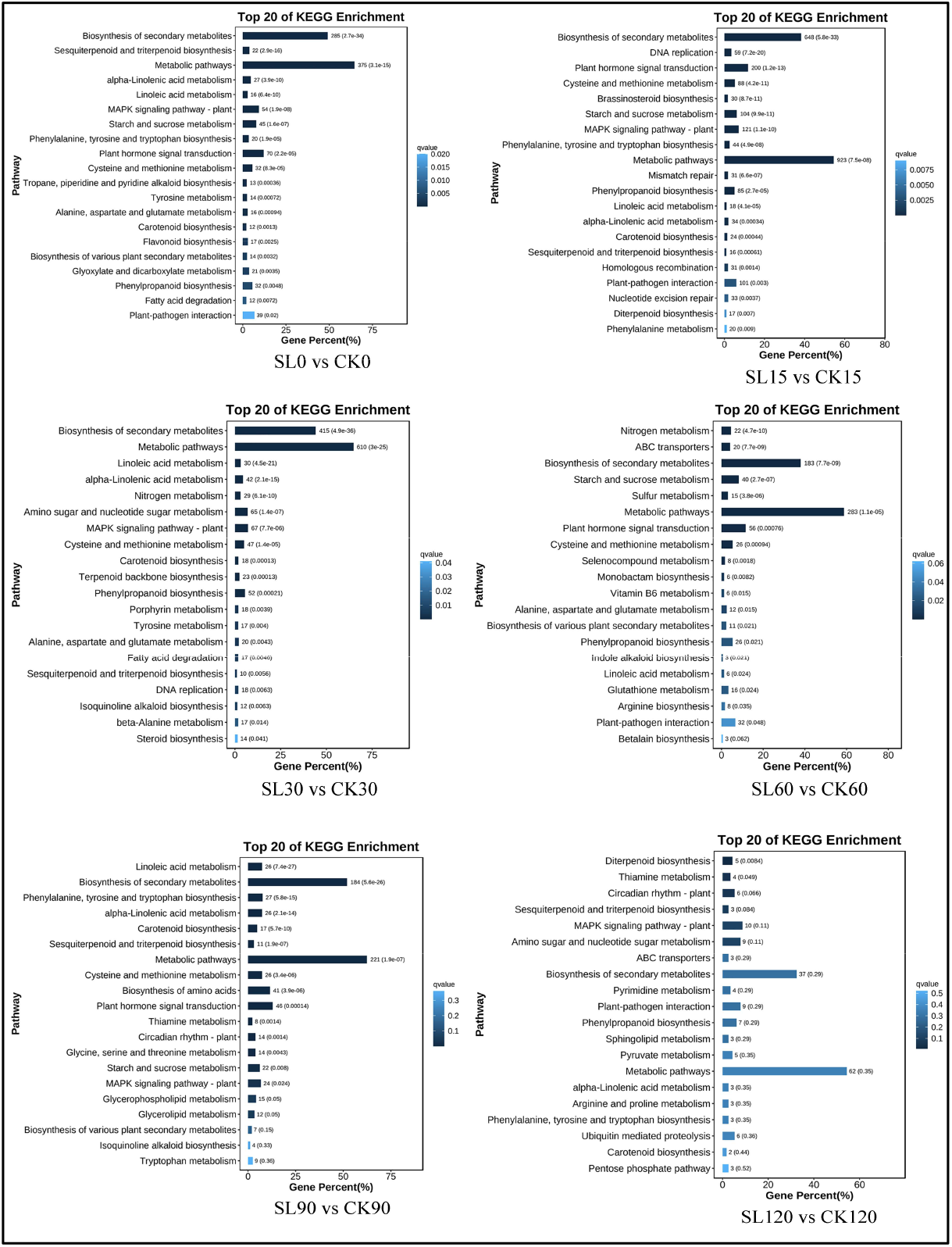
KEGG enrichment analysis of differentially expressed genes in different periods

### WGCNA Co-expression Network Analysis

To elucidate the regulatory network of strigolactones (SLs) in Pisang Awak banana corm development, a weighted gene co-expression network (WGCNA) was constructed based on transcriptome data, dividing genes into multiple modules (Figure 6). Correlation analysis between module eigengenes (MEs) and corm diameter traits identified three modules— greenyellow, Mediumpurple2, and coral—that were significantly associated with corm weight, height, and diameter (Figure 7). Among them, only the greenyellow module contained the D53 gene, a key component of the SL signaling pathway, and showed the highest correlation with corm weight, indicating that it is likely the core module directly regulated by SLs. Cytoscape visualization revealed that within this module, SPL15 acted as a central hub, highly connected to genes involved in multiple hormone signaling pathways, including auxin (IAA), cytokinin (CTK), abscisic acid (ABA), gibberellins (GA), brassinosteroids (BRs), and jasmonic acid (JA). These findings suggest that SL signaling regulates SPL15 via D53, which in turn integrates multiple hormone pathways to control corm development, forming a core regulatory network: SL → D53 → SPL15 → other hormone pathway genes (Figure 8). This result further confirms the central role of SPL15 in SL-mediated regulation and hormone crosstalk and provides candidate genes and mechanistic insights for subsequent functional validation.

**Fig. 6.**
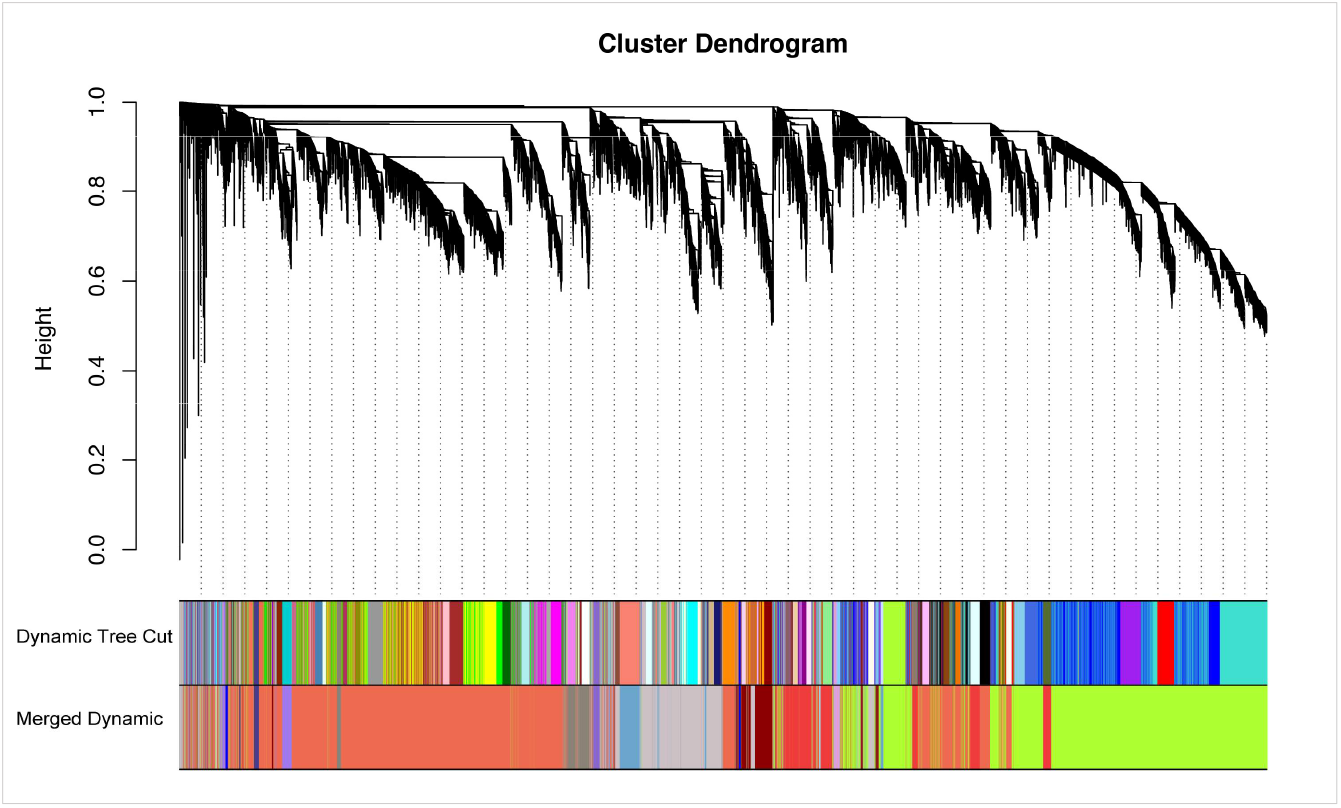
Gene dendrogram after dynamic tree cutting and after dynamic merging.

**Fig. 7.**
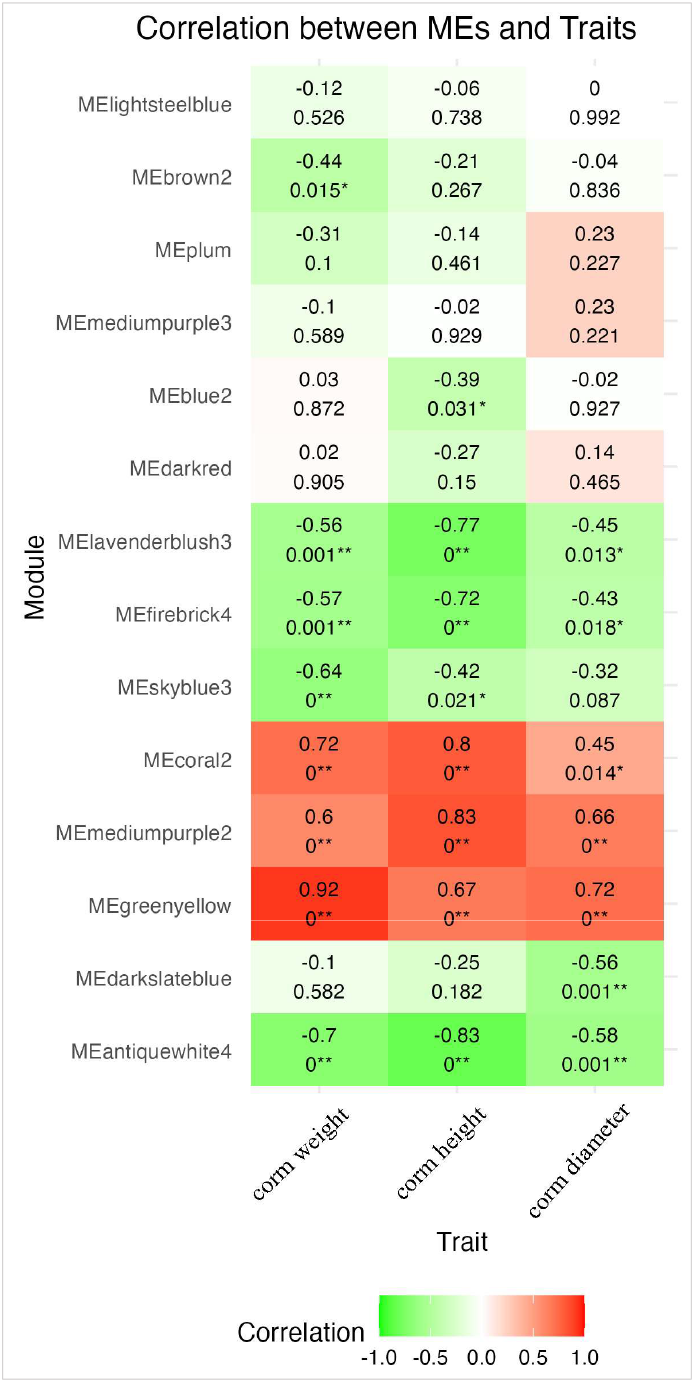
Correlation of each module identified by WGCNA to hormones and height. The upper number in each small square represents the Pearson correlation coefficient, and the number in parentheses represents the significance (P-value) of each module compared to each external factor.

**Fig. 8.**
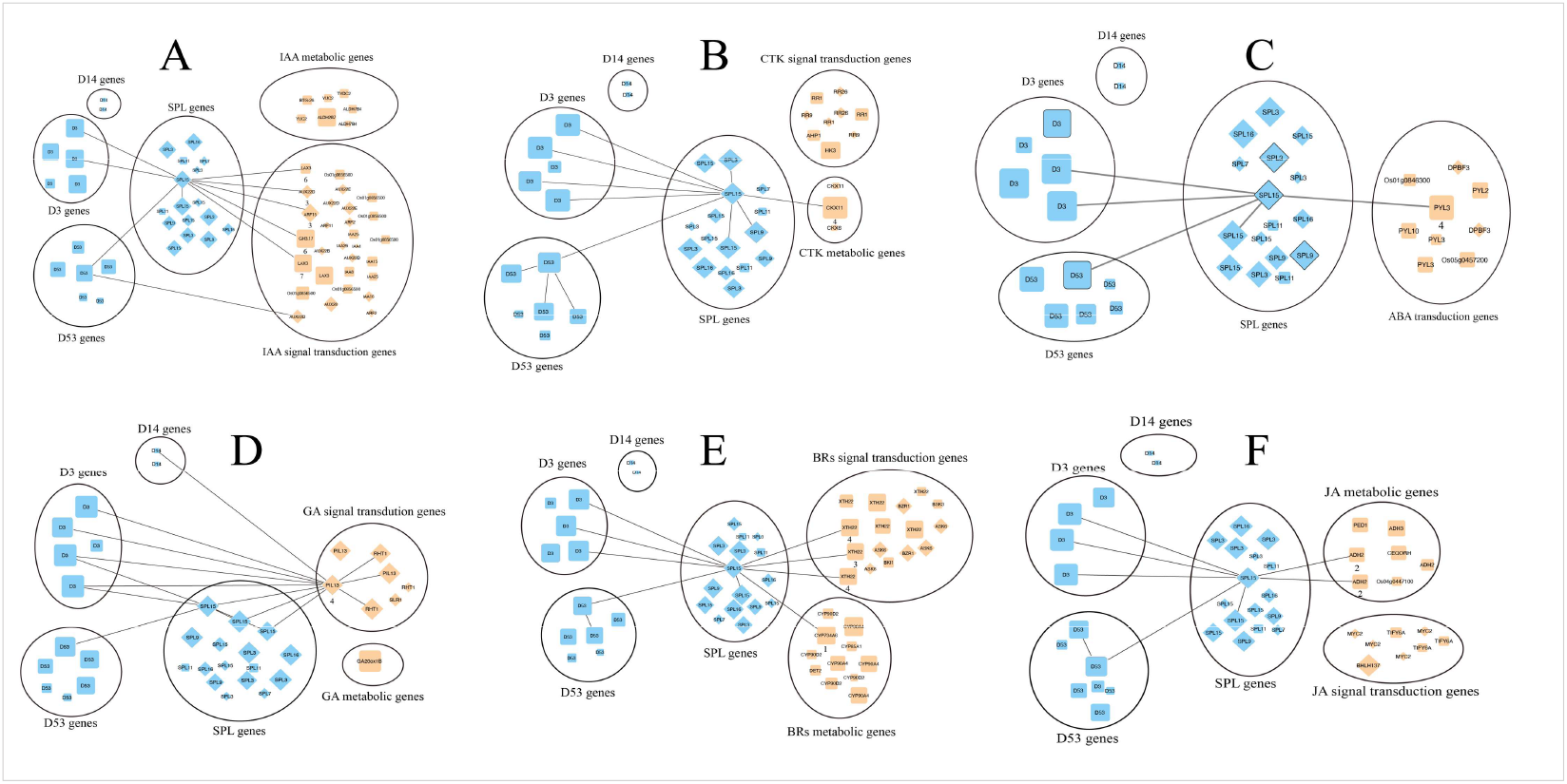
SL → D53 → SPL15 Regulatory Network Revealed by WGCNA in Pisang Awak banana Corm Development. Numbers below the color blocks represent the number of SPL motifs (GTAC).

### Real-time quantitative PCR validation results

To validate the reliability of the transcriptome sequencing data, seven key genes associated with SL regulation and hormone crosstalk, including SPL15 and other hormone pathway genes directly connected to SPL15 within the regulatory network, were analyzed by qRT-PCR. The results demonstrated that the relative expression levels of these genes across different time points were highly consistent with the RNA-seq data. Correlation analysis between qRT-PCR and RNA-seq expression levels yielded R^2^ = 0.9915, indicating a strong concordance between the two datasets (Figure 9). These findings confirm the accuracy and reliability of the transcriptome data and further validate the importance of the selected genes within the SL-mediated hormone crosstalk network, providing a solid foundation for subsequent functional analyses.

**Fig. 9.**
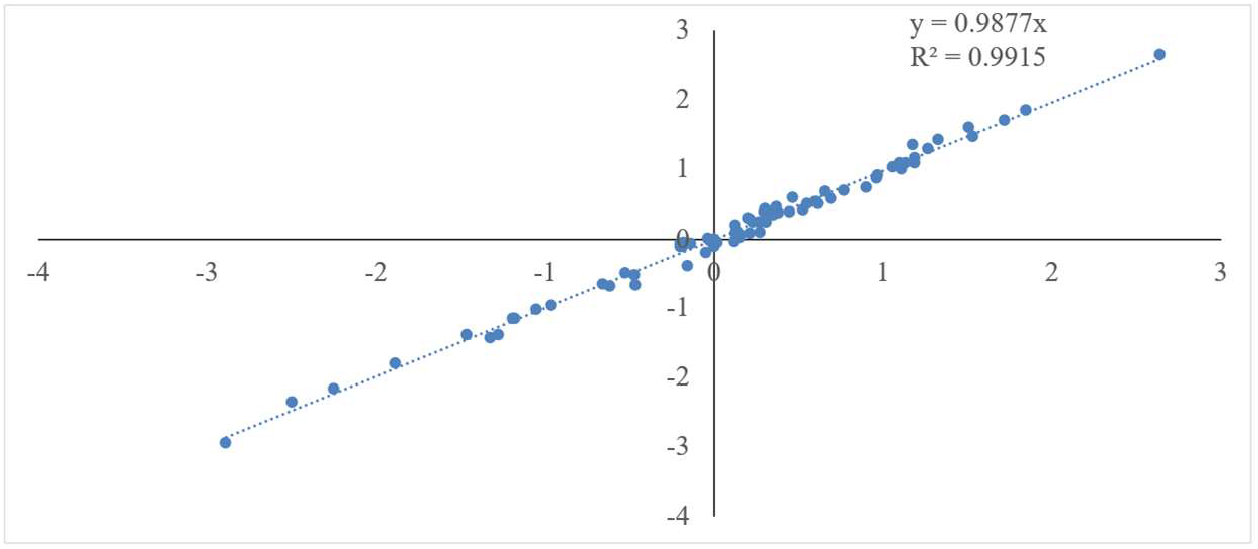
Correlation analysis between qRT-PCR and transcriptome data. The x-axis represents log_2_ fold change, and the y-axis represents -ΔΔCt.

## Discussion

The interactions among various plant hormones are commonly referred to as “crosstalk.” Strigolactones (SLs) engage in multiple direct and indirect interactions with other hormones ^13^. To date, studies on SL crosstalk with other plant hormones have mainly focused on the regulation of lateral bud and root development ^14^. In the present study, banana corms, a type of modified stem, were used for the first time as the research subject to investigate the mechanisms underlying SL-mediated crosstalk with other plant hormones during corm growth.

Banana corms are modified stems whose growth and development are regulated by multiple hormones. Previous studies have shown that strigolactones (SLs) exhibit concentration-dependent effects on stem growth: low concentrations of SLs generally promote stem elongation and development, whereas high concentrations inhibit growth. For example, in cucumber, treatment of ‘LZ1’ seedlings with 1 μmol·L^−1^ and 5 μmol·L^−1^ GR24 increased average plant height by 19.55% and 4.32% and stem diameter by 14.93% and 11.57% compared with the control, respectively. In contrast, treatment with 10 μmol·L^−1^ GR24 significantly reduced average plant height by 33.40% and stem diameter by 16.92% relative to the control ^15^. In this study, a relatively high concentration of 30 μmol·L^−1^ rac-GR24 was applied, resulting in significant reductions in corm weight, height, and diameter, indicating an inhibitory effect (Figure 1). This inhibition differs from the SL-mediated regulation of axillary buds, where increasing SL concentrations do not promote lateral branch development. These findings suggest that SLs exert organ-specific effects, with differential responses in corms and axillary buds.

Strigolactones (SLs) are pivotal phytohormones regulating plant architecture, and their canonical signal perception and transduction mechanisms have been extensively characterized. Previous studies have shown that SL molecules are perceived by the receptor protein D14, which recruits the F-box protein MAX2/D3, thereby promoting ubiquitination and proteasomal degradation of the core repressor SMXLs/D53. This degradation relieves D53-mediated transcriptional repression of downstream genes, triggering a signaling cascade that remodels plant development ^16^. Within this core signaling axis, the SQUAMOSA promoter binding protein-like (SPL) transcription factor family has been identified as a key downstream effector of SL signaling. In model plants such as rice, degradation of D53 releases the transcriptional activity of SPL proteins, such as OsSPL14/IPA1, which subsequently suppress lateral bud elongation by mediating local cytokinin (CTK) degradation ^17, 18^. However, the systemic regulatory role of SPL transcription factors in the development of subterranean modified stems, such as banana corms, remains largely unexplored.

In the present study, through WGCNA and targeted co-expression network analysis, we not only confirmed a strong activation of the D53-SPL core signaling module in the corm apical meristem but also made a groundbreaking discovery: SPL functions not merely in a single downstream pathway but as a super “hub,” forming highly connected co-expression relationships with nearly all major plant hormone signaling and metabolic pathway genes. Our network topology clearly demonstrates that the D53-SPL module extensively links key genes across multiple hormone pathways, including those involved in auxin polar transport and response (LAX3, AUX22D, ARF15), gibberellin (GA) and brassinosteroid (BR)-mediated cell elongation (PIL13, XTH22, CYP734A6), cytokinin degradation (CKX11), and ABA/JA-mediated metabolic homeostasis (PYL3, ADH2). These results indicate that, during SL-induced morphological remodeling of banana corms—characterized by increased longitudinal growth and reduced lateral diameter—the D53-SPL module acts as a global orchestrator of hormone crosstalk. It not only mediates individual hormone responses but also integrates the spatiotemporal distribution and signaling intensity of multiple hormones such as IAA, GA, BR, and CTK, thereby disrupting the original developmental equilibrium of the corm and precisely driving organ elongation and resource accumulation. This finding not only extends our understanding of SL signaling within multi-hormone regulatory networks but also provides a new theoretical framework for elucidating the molecular basis of banana corm development and guiding future crop improvement.

